# Stem cell-free therapy for glaucoma to preserve vision

**DOI:** 10.1101/2021.06.18.449038

**Authors:** Ajay Kumar, Xiong Siqi, Minwen Zhou, Wen Chen, Enzhi Yang, Andrew Price, Liang Le, Ying Zhang, Laurence Florens, Michael Washburn, Akshay Kumar, Yunshu Li, Yi Xu, Kira Lathrop, Katherine Davoli, Yuanyuan Chen, Joel S. Schuman, Ting Xie, Yiqin Du

**Affiliations:** Department of Ophthalmology, University of Pittsburgh, Pittsburgh, PA 15213; Stowers Institute of Biomedical Research, Kansas City, Missouri 64110; Department of Cardiothoracic Surgery, University of Pittsburgh, Pittsburgh, PA 15213; McGowan Institute for Regenerative Medicine, University of Pittsburgh, Pittsburgh, PA 15213; Department of Ophthalmology, New York University School of Medicine, New York, NY, USA 10017; Department of Developmental Biology, University of Pittsburgh, Pittsburgh, PA 15213

## Abstract

Glaucoma is the leading cause of irreversible blindness with trabecular meshwork (TM) dysfunction resulting in elevated intraocular pressure and retinal ganglion cell (RGC) damage leading to vision loss. In this study, we discovered that secretome, derived from human TM stem cells, via minimal invasive periocular injection, can reduce intraocular pressure, restore TM homeostasis, protect RGC, and restore RGC function in both steroid-induced and genetic myocilin mutant mouse models of glaucoma. The secretome upregulated the COX2-PGE2 axis via mitochondrial TMEM177 and led to activation of endogenous stem cells and TM regeneration. Inhibition of COX2 abolished the protective effect of secretome on TM cells. Secretome treatment also enhanced RGC survival and function. Proteomic analysis revealed that the secretome is enriched with proteins involved in extracellular matrix modulation leading to the remodeling of TM to restore homeostasis. This study highlights the feasibility of stem cell-free therapy for glaucoma with minimal invasive administration and the involvement of multiple novel pathways for a cumulative regenerative effect on the TM to protect RGC.

**Brief summary:** This study describes a cell-free treatment using stem cell secretome in two animal models of glaucoma and explores the potential mechanisms

## Introduction

Glaucoma is the leading cause of irreversible blindness and associated with huge socioeconomic burden^1^. Glaucoma is one of the most prevalent visual disorders affecting 76 million people worldwide in 2020 and estimated to affect 111.8 million by 2040^2^. In glaucoma, the trabecular meshwork (TM) structure and function is abnormal and the retinal ganglion cells (RGC) are degenerated^3^. The TM tissue together with the Schlemm’s canal endothelium provides resistance to aqueous humor outflow. TM cellularity reduces with age at an average rate of 0.58% per year^4^ and the reduction is further aggravated in glaucoma resulting in pathological conditions including trabecular fusions and abnormal extracellular matrix (ECM) deposition, causing increased outflow resistance and elevated intraocular pressure (IOP)^5,6^. RGC death in glaucoma is caused by elevated IOP, hypoxia, increased reactive oxygen species and T-cell infiltration, and enhanced production of inflammatory cytokines^7^.

Reducing IOP is currently the only treatment target for glaucoma, including eye drops and medications (such as beta blockers, alpha agonists, carbonic anhydrase inhibitors, Rho kinase inhibitors, and prostaglandin analogs), laser procedures, and surgeries. These treatments are effective but have their limitations and side effects and do not target the TM pathophysiologic change which is the main reason of IOP elevation. Hence new therapies targeting the TM with long-term efficacy are needed for glaucoma. TM stem cells (TMSC)^8-10^, mesenchymal stem cells^11,12^, adipose-derived stem cells^13^, and induced pluripotent stem cells (iPSC)^14^ have shown promising effects on regenerating the TM and preventing vision loss.

Secretome is secreted by cells including cytokines, growth factors, extracellular matrix proteins, microRNAs, non-coding RNAs, and mRNAs. Stem cell secretome, exosomes, and microvesicles have shown therapeutic value in various disorders^15-18^. However, no studies have investigated the roles of stem cell secretome on TM regeneration for glaucoma treatment and the mechanisms of secretome-induced regeneration remain elusive.

Our recent work has demonstrated that TMSC can impart therapeutic effects in glaucoma^9,10^. In this study, we explored the therapeutic effects of secretome derived from human TMSC (TMSC-Scr) with minimal invasive periocular injection in two mouse models of glaucoma: an ocular hypertension model induced by dexamethasone-acetate (Dex-Ac) periocular injection^19^ and a transgenic open-angle glaucoma model with myocilin Y437H mutation (Tg-MyocY437H)^20^, and uncovered the potential mechanisms. We explored that the mice received TMSC-Scr treatment increased the TM cellularity, reconstructed the TM ECM, reduced IOP, prevented/rescued RGC death, and preserved/rescued RGC function. We discovered that endogenous TMSC were activated via upregulation of the cyclooxygenase-2 and prostaglandin E2 (COX2-PGE2) signaling axis. Proteomic analysis indicates that TMSC-Scr is enriched with proteins involved in ECM modulation which are associated with TM ECM remodeling. Our study provides the first proof of concept for the use of stem cell secretome for glaucoma treatment and uncovers the potential mechanisms involved.

## Results

### TMSC secretome prevents dexamethasone-induced glaucomatous changes in TM cells *in vitro*

We cultured TMSC from human donor eyes as previously reported^21,22^ and characterized each TMSC strain from different donors by flow cytometry showing positive expression of stem cell markers CD90, CD73, CD105, CD166, SSEA4, OCT4, ABCG2, STRO-1, NOTCH-1, CD271, and negative expression of CD34 and CD45 (**Supplementary Fig. 1a-b**). This demonstrates the stem cell nature of human TMSC used in this study, similar to what we previously reported^21,22^. To ensure secretome was harvested from healthy cells after 48h of starvation, we stained the cells with Annexin V and 7-Aminoactinomycin D (7-AAD) **(Supplementary Fig. 1c**) and with Calcein AM/Hoechst 33342 (**Supplementary Fig. 1d**), which showed >95% viable cells after secretome harvesting. The results confirmed reasonably pure secretome released by live cells with less than 5% cell death as required^23^. Secretome from human corneal fibroblasts^8,24^ served as a control and the viability of fibroblasts was also confirmed **(Supplementary Fig. 1c-d)**. To evaluate if TMSC secretome was cytotoxic, we treated human TM cells with TMSC-Scr for 48h and did MTT and alamarBlue assays. The results showed no significant difference in cell viability and proliferation between cells with and without TMSC-Scr treatment **(Supplementary Fig. 1e-f)**. Collectively, these assays demonstrate that the secretome was harvested from healthy cells and had no cytotoxicity to human TM cells, supporting its safety.

Then we examined if TMSC secretome has protective roles to human TM cells *in vitro*. One of the characteristics of TM cells is that they are responsive to dexamethasone (Dex) treatment with increased intracellular myocilin expression^10,25,26^. Myocilin^27,28^ and angiopoietin-like 7 (ANGPTL7)^22,29^ are both glaucoma-associated markers. Chitinase 3 Like 1 (CHI3L1) is involved in ECM remodeling in the outflow pathway and has been used as a TM cell marker as has water channel protein aquaporin 1 (AQP1)^21,30^. Fibronectin, an ECM component in the TM, is increased in glaucoma and is believed to contribute to the TM stiffness and increased outflow resistance^31^. When cultured human TM cells were treated with 100 nM Dex for 5 days, the TM cells had reduced expression of CHI3L1 and AQP1 and increased expression of myocilin, ANGPTL7, and fibronectin as compared to no-Dex control (**Supplementary Fig. 2a-d**). To assess TMSC-Scr therapeutic effect, TMSC-Scr was added together with Dex for 5 days (Parallel TMSC-Scr, **Supplementary Fig. 2a**) for prevention effect assessment, or together with Dex for another 5-day after the initial 5-day Dex-alone treatment (Post TMSC-Scr, **Supplementary Fig. 2b**) for reversal effect assessment. TMSC-Scr prevented the reduction as well as restored the expression levels of TM cell markers CHI3L1 and AQP1, and prevented as well as reduced myocilin, ANGPTL7, and fibronectin expression as demonstrated by immunofluorescent staining (**Supplementary Fig. 2a-g)**. The mRNA levels of *MYOCILIN* (**Supplementary Fig. 2h**) and *ANGPTL7* (**Supplementary Fig. 2i**) were dramatically reduced by both parallel- and post-TMSC-Scr treatments. These results indicate that TMSC-Scr can prevent and reverse steroid-induced glaucomatous changes in cultured human TM cells.

### TMSC secretome reduces IOP and prevents RGC loss in steroid-induced ocular hypertension mouse model

To induce an ocular hypertension model, we periocularly injected 200 µg/20 µl of Dex-Ac into adult C57BL/6J mice once a week for 6 weeks and sacrificed the animals at week-8. The mouse IOP started to elevate from week-1 after Dex-Ac injection and remained elevated up to week-8 after Dex-Ac injection had been stopped for two weeks (**Fig. 1a-b**). 20 µl of 25x concentrated TMSC-Scr or fibroblast-secretome (Fibro-Scr) was injected periocularly once a week starting at week-3 and ending at week-6. The injections of Dex-Ac and secretome from week-3 to week-6 were given at the same time but at different periocular regions (superior and inferior) to avoid reagents being intermixing before getting into target sites. One week after the first TMSC-Scr injection, the IOP reduced to 13.8±0.5 mmHg (week-4) as compared to mice treated with Dex-Ac alone (15.9±0.4 mmHg) and then reduced to normal range from week-5 to -8 (10.7±0.4 mmHg), similar to that of vehicle control (12.2±0.6 mmHg). However, Fibro-Scr did not reduce the Dex-Ac elevated IOP (16.2±0.8 mmHg at week-4 and 15.4±0.5 mmHg at week-8) but remained similar to Dex-Ac treated eyes (15.9±0.9 mmHg at week-4, 15.2±0.3 mmHg at week-8). The TM cells on plastic sections obtained at week-8 were counted with the results shown in **Fig. 1c-d**: TMSC-Scr injection increased the TM cellularity (19.1±1.7 cells/TM section) as compared to Dex-Ac mice (14.7±1.3), similar to normal control (20.7±1.5), while Fibro-Scr treatment could not increase TM cell number significantly (16.7±1.3). There was a significant increase of myocilin protein level in the limbal tissue (including the TM) of Dex-Ac mice detected by immunoblotting, which was reduced after TMSC-Scr treatment **(Fig. 1e)**. Immunoblotting on the aqueous humor revealed higher level of secreted myocilin in TMSC-Scr treatment group than that in Dex-Ac mice **(Fig. 1f)**.

**Fig. 1.**
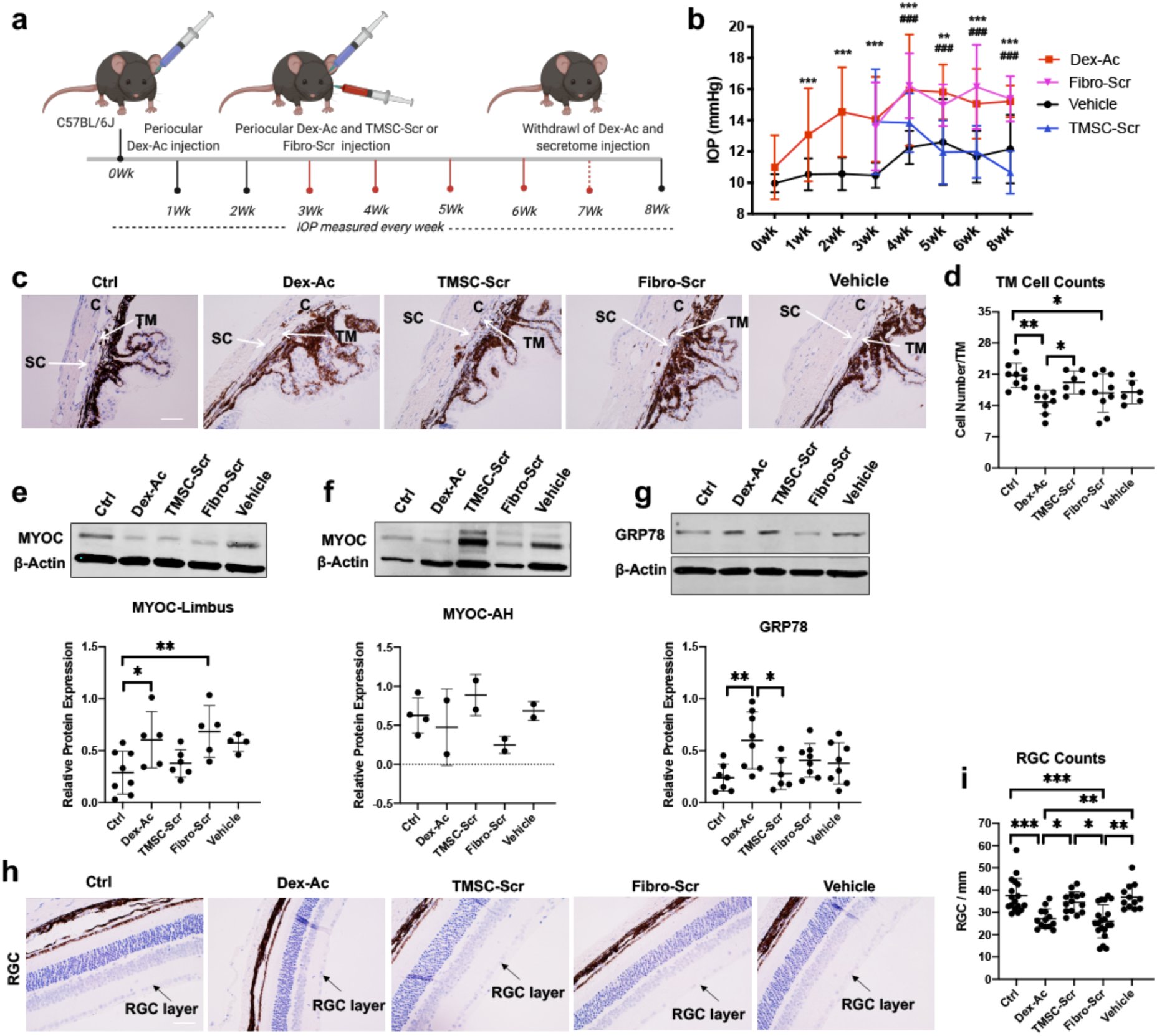
TMSC secretome reduces IOP, restores TM cellularity, and modulates myocilin in Dex-Ac mice. **a**. Schematic showing procedures for Dex-Ac and secretome treatment in Dex-Ac induced mouse model, **b**. IOP dynamics in Dex-Ac induced model showing higher IOP in Dex-Ac and fibro-Scr treated animals while treatment with TMSC-Scr reduced the IOP to normal level. *comparison between vehicle and Dex-Ac mice, #comparison between Dex-Ac and TMSC-Scr treated mice, (n=10-21), ***/###p<0.001, Mean±SEM, Two-way ANOVA followed by Tukey posttest. **c**. DIC monocolor images showing anterior angle structure in Dex-Ac mice, **d**. Bar diagram showing quantification of TM cell number in Dex-Ac mice (n=3-4), **e-f**. Immunoblotting bands and bar diagram showing protein expression and quantification for Myoc in the limbus tissue and aqueous humor respectively. **g**. Immunoblotting expression and quantification of GRP78 in limbus tissue, limbus (n=4), AH (n=2), **h**. DIC monocolor images showing RGC layer in Dex-Ac mice, **i**. Bar diagram showing quantification of RGC number, (n=3-4). Multiple dots in bar graphs represent combined results of biological and technical replicates, scale bar-100 µm. **d-i**, *P<0.05, **P<0.01. Mean±SD, One-way ANOVA followed by Tukey posttest. SC-Schlemm’s canal, TM-Trabecular Meshwork, C-Cornea.

Cells develop endoplasmic reticulum (ER) stress when the ER is burdened by increased accumulation of misfolded/unfolded proteins. Increased activation of chronic ER stress has been observed in the TM of human and mouse glaucoma^32^, which is responsible for enhanced cell death. A significantly increased expression of GRP78, an ER stress^33,34^, in the limbal tissue in Dex-Ac mice was detected as compared to the control, while the GRP78 expression was reduced after TMSC-Scr treatment **(Fig. 1g)**. Although, RGC can regenerate in embryo and neonates, this ability is diminished few days after birth due to intrinsically controlled molecular mechanisms^35^. Dex-Ac mice at week-8 showed significant RGC loss (28.4±5.4/mm) as compared to the WT control (37.6±7.6/mm) and vehicle control (36.9±5.6/mm) as counted on the plastic sections of the mouse eyes at week-8, while the RGC loss was prevented with TMSC-Scr treatment (34.5±4.7/mm) **(Fig. 1h-i)**. The results indicate that TMSC-Scr promotes myocilin secretion and modulates Dex-induced ER stress of the TM cells, which contributes to the increased TM cellularity and reduced IOP. TMSC-Scr also prevents RGC loss in the steroid-induced mouse model.

### TMSC secretome activates the COX2-PGE2 pathway to activate endogenous stem cells

COX2 (cyclooxygenase) is a major protein that helps the biogenesis of prostaglandin E2 (PGE2) from prostaglandin H2 (PGH2) in response to physiological demand^36^. The presence of PGE2 in TM cells plays an important role in maintenance of IOP in normal range^37^. Human TM cells secrete PGE2 which is abrogated by glucocorticoid treatment^38^. Dex treatment dramatic reduced COX2 expression in cultured TM cells *in vitro* which was restored by TMSC-Scr treatment **(Fig. 2a)**. Transmembrane protein 177 (TMEM177), a mitochondrial protein, acts upstream of COX2 to increase and stabilize COX2 expression^39^. Analysis of TMEM177 in cultured human TM cells showed a diminished expression after Dex treatment, which was restored after parallel- and post-TMSC-Scr treatments (**Fig. 2b)**. Immunofluorescent staining and immunoblotting of mouse limbal tissue showed that COX2 levels were reduced in Dex-Ac treated tissue and increased after TMSC-Scr treatment at week-8 **(Fig. 2c-d**). Immunoblotting on the mouse limbal tissue showed that TMEM177 levels had similar changes to that of COX2 (**Fig. 2e)**. PGE2 secretion to the mouse aqueous humor was detected and compared by ELISA (**Fig. 2f**). PGE2 secretion was reduced after Dex-Ac treatment and was restored to normal level after TMSC-Scr treatment, but not after Fibro-Scr treatment. PGE2 has been reported to have the ability to maintain self-renewal of mesenchymal stem cells^40^. Here, we observed that the ABCB5^+^ and OCT4^+^ stem cell population as well as Ki67^+^ proliferative cells in mouse limbus and TM tissue were diminished after Dex-Ac treatment and increased after TMSC-Scr treatment **(Fig. 2g-h)**. These results indicate that the TMSC-Scr is capable of activating COX2-PGE2 signaling to sustain TMSC and TM cells for steroid-induced glaucoma.

**Fig. 2.**
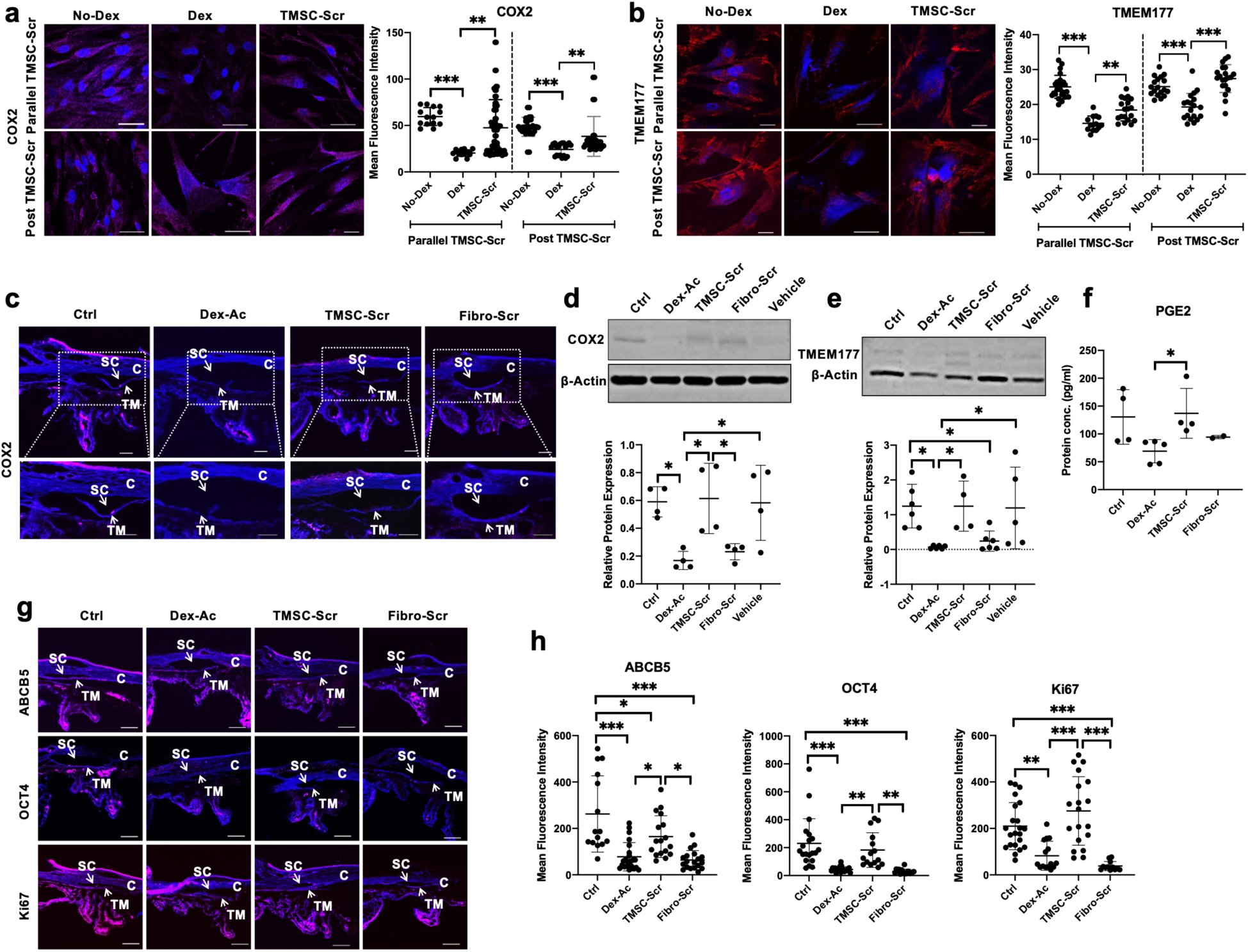
TMSC secretome modulates COX2-PGE2 signaling in Dex-Ac mice. **a**. Immunofluorescent staining and quantification of COX2 in TMSC-Scr (four different TMSC-Scr) treated cells (three different TM cell strains) in parallel and post Dex induction. Multiple dots in bar graphs represent combined results of biological and technical replicates, **b**. Protein expression profile showing immunofluorescent staining and quantification of TMEM177 in secretome treated cells in parallel and post Dex induction, **c**. Immunofluorescent images showing protein expression of COX2 in limbus tissue of Dex-Ac mice, **d-e**. Immunoblotting showing protein expression profile and quantification of COX2 and TMEM177 in limbus tissue of, (n=4). Multiple dots in bar graphs represent combined results of biological and technical replicates, **f**. Bar diagrams showing levels of PGE2 in aqueous humor secreted in different treatment groups of Dex-Ac mice (n=4-6), **g-h**. Immunofluorescent images and bar graphs showing protein expression and quantification of ABCB5, OCT4 and Ki67 in Dex-Ac mice, (n=3), multiple dots on bar graph indicates fluorescence intensity values measured from at least 3-16 different limbus sections of the eye. scale bar-50 um. *p<0.05, **p<0.001, ***p<0.0001. Mean±SD, One-way ANOVA followed by Tukey posttest. SC-Schlemm’s canal, TM-Trabecular Meshwork, C-Cornea.

### TMSC secretome restores TM cellularity and reduces IOP in Tg-MyocY437H mice

To further confirm the TMSC-Scr therapeutic effects, we also investigated a genetic mouse model of glaucoma, the Tg-MyocY437H mice^10,20^, which start to have elevated IOP at 3-4-month of age. We periocularly injected 20 μl of 25x concentrated TMSC-Scr or 20 μl plain medium as sham control into the Tg-MyocY437H mice at 4-month of age (week-0) once a week until week-7 and sacrificed the mice at week-10 (**Fig. 3a**). IOP reduction started at 1-week after TMSC-Scr treatment and remained reduced until week-10 as compared to Tg-MyocY437H mice without any treatment (Tg-Myoc) and with medium injection (sham), and similar to wildtype (WT) littermates (**Fig. 3b**). Tg-MyocY437H mice (14.8±2.5 mmHg) and sham control (15.0±1.7 mmHg) still had elevated IOP at week-10, whereas TMSC-Scr treated mice maintained the IOP at normal range (9.5±2.2 mmHg), comparable to WT control (10.3±2.2 mmHg). The TM cells on plastic sections were counted shown that Tg-MyocY437H mice had reduced number of TM cells (10±1.4/TM section) as compared to WT mice (16.4±2.1/TM) at week-10 (**Fig. 3c-d)**. Similar to its effects to the Dex-Ac mice, treatment with TMSC-Scr significantly increased the TM cellularity in Tg-MyocY437H mice (17.5±1.1/TM). One feature of the Tg-MyocY437H mice is that mutant Myoc cannot be secreted out but stuck in the ER of TM cells leading to ER stress^10,20^. Indeed, we found an increased level of Myoc accumulated in the limbus tissue of the Tg-MyocY437H mice, which was reduced to the level as WT control after TMSC-Scr treatment as detected by staining **(Fig. 3e)** and by immunoblotting **(Fig. 3f)**. TMSC-Scr treatment also led to increased secretion of Myoc into the aqueous humor **(Fig. 3g)**. Similar to Dex-Ac mice, GRP78 expression was significantly increased in Tg-MyocY437H mice and reduced to WT level after TMSC-Scr treatment **(Fig. 3h)**. Using an anti-CD31 antibody to stain the Schlemm’s canal and vascular endothelium to mark the TM tissue location^10,14,26^, we observed an increased expression of ECM markers fibronectin and collagen IV in the TM in Tg-MyocY437H mice, which was reduced after TMSC-Scr treatment while sham group showed no reduction **(Supplementary Fig. 3a-b)**. Similar to Dex-Ac mice, an analysis of COX2-PGE2 signaling axis in Tg-MyocY437H mice showed increased levels of COX2 and TMEM177 by immunoblotting (**Fig. 3i-j**), PGE2 by ELISA (**Fig. 3k**), ABCB5^+^ and OCT4^+^ stem cells, and Ki67^+^ dividing cells by immunostaining (**Supplementary Fig. 4a-d**) after TMSC-Scr treatment. These results further confirmed the therapeutic effects of the TMSC-Scr in the Tg-MyocY437H mouse model of glaucoma.

**Fig. 3.**
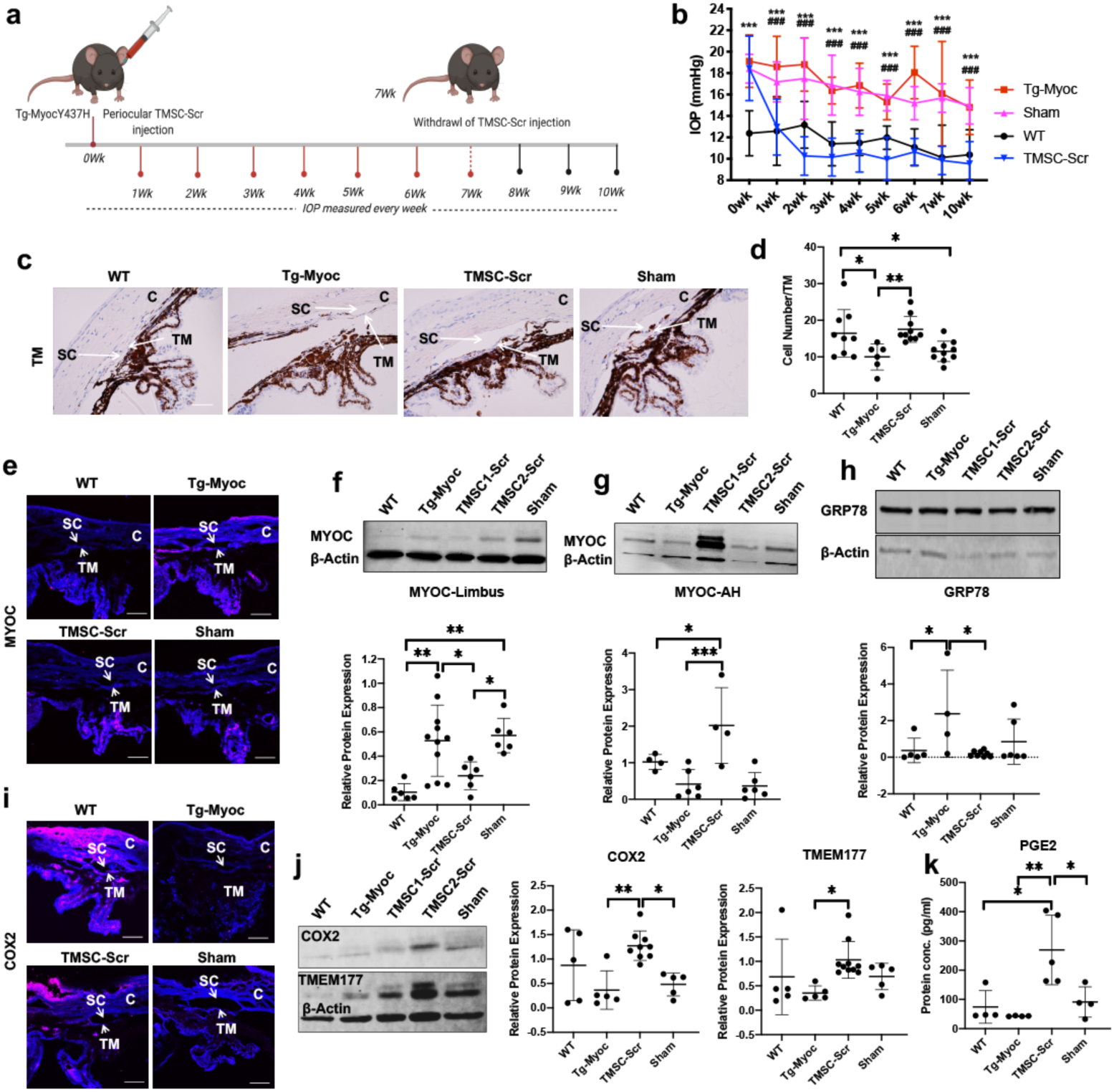
TMSC secretome reduces IOP, restore TM homeostasis and modulates COX2-PGE2 signaling in Tg-MyocY437H mice. **a**. Schematics showing procedure followed for TMSC-Scr treatment in Tg-Myocy437H mice, **b**. IOP levels in Tg-MyocY437H mice showing IOP reduction by TMSC-Scr while no effect on sham treated Tg-MyocY437H mice. *comparison between WT and Tg-MyocY437H mice, #comparison between Tg-MyocY437H and TMSC-Scr treated mice, (n=10-32), ***/###p<0.001, Mean±SEM, Two-way ANOVA followed by Tukey posttest. **c**. DIC monocolor images showing TM cells in trabecular meshwork in Tg-MyocY437H mice, scale bar-100 um, **d**. Bar diagram showing quantification of TM cell number in Tg-MyocY437H mice (n=3-4), **e**. Immunofluorescent images showing protein expression of MYOC in limbus tissue of Tg-MyocY437H mice, **f-h**. Immunoblotting showing protein expression. and quantification of of Myoc in limbus tissue and aqueous humor respectively, and that of GRP78 in limbus of different treatment groups in Tg-MyocY437H mice respectively, limbus (n=4). Multiple dots in bar graphs represent combined results of biological and technical replicates, AH (n=4-6), **i**. Immunofluorescent images showing protein expression of COX2 in limbus tissue of Tg-MyocY437H mice, SC-Schlemm’s canal, TM-Trabecular Meshwork, **j**. Immunoblotting showing protein expression and quantification profile of COX2 and TMEM177 in limbus tissue, (n=4). Multiple dots in bar graphs represent combined results of biological and technical replicates, **k**. Bar diagrams showing levels of PGE2 in aqueous humor secreted in different treatment groups of Tg-MyocY437H mice (n=4-6).

Since there is a dramatic reduction in COX2-PGE2 signaling axis in the glaucomatous TM tissue of both Dex-Ac and Tg-MyocY437H mice, we investigated whether COX2 is altered in the human glaucomatous TM tissue. Immunostaining shows that both COX2 and TMEM177 expression was dramatically reduced in the human glaucomatous TM tissue (**Supplementary Fig. 5a-b**). Interestingly, the expression of ABCB5 and OCT4 was also reduced in the human TM tissue indicating a reduction of endogenous TMSC in glaucomatous TM tissue (**Supplementary Fig. 5a-b**).

Both nimesulide^41^ and indomethacin^42^ have been reported to inhibit COX2. We abrogated the expression of COX2 in primary human TM cells by treatment of optimized dosage of nimesulide (200µM) and indomethacin (100µM). Both the inhibitors reduced COX2 expression dramatically after 10 days of treatment in both control TM cells and those treated with TMSC-Scr, resulting in the loss of protective effect of TMSC-Scr on TM cell function, as evidenced by reduced expression of CHI3L1 and increased expression of myocilin (**Fig. 4a-d**).

**Fig. 4.**
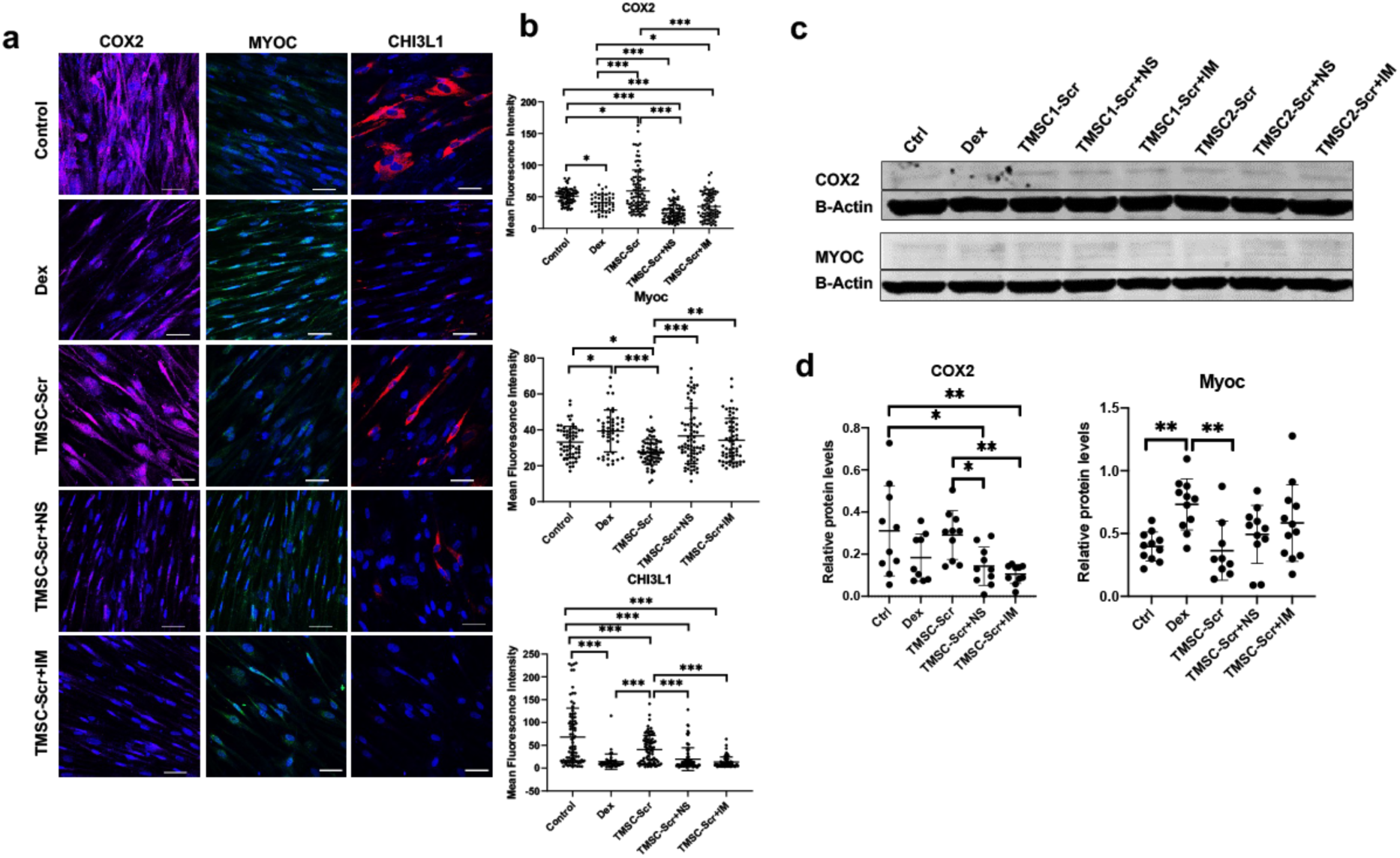
Inhibition of COX2 leads to abolishment of protective effect of secretome on TM cells. **a**. Immunofluorescent images showing effect of inhibitor (Nimesulide, NS and Indomethacin, IM) treatment on staining patterns of COX2, MYOC, and CHI3L1, scale bar-50 µm, **b**. Bar graphs showing quantification of protein expression for COX2, MYOC, and CHI3L1 after inhibitor treatment, n=3 different TM cell strains were treated with two different TMSC secretomes. Multiple dots in bar graphs represent combined results of individual cells quantified in terms of fluorescence in different groups, **c-d**. Immunoblot and bar graphs showing quantification of protein expression of COX2 and Myocilin after inhibitor treatment. Multiple dots in bar graphs represent combined results of biological and technical replicates. *P<0.05, **p<0.001, ***P<0.0001. Mean±SD, One-way ANOVA followed by Tukey posttest.

### TMSC secretome prevents RGC death *in vitro* and *in vivo*

Preserving and restoring the RGC and their function is the ultimate goal of glaucoma treatment. Glaucoma progression is accompanied by increased loss of RGC at structural and functional level^43^. Pattern electroretinography (PERG) is an optimal approach to evaluate RGC function^10,44^. PERG shows that TMSC-Scr treatment preserved the RGC function of Tg-MyocY437H mice as indicated by increased amplitude of P1 wave (7.74±1.75 µV), similar to that of WT (7.76±1.14 µV), while the sham treatment could not recover P1 amplitude (4.05±1.95 µV), similar to that of untreated Tg-MyocY437H (5.86±2.24 µV) (**Fig. 5a-b**). RGC numbers counted from retinal plastic sections showed an average of about 36% loss of the RGC of Tg-MyocY437H mice (24.3±7.8 cells/mm) as compared to WT (37.8±9.3 cells/mm). This loss was rescued by TMSC-Scr treatment (37.3±9.4 cells/mm) while no protective effect was observed in the sham group (28.5±9.2 cells/mm) (**Fig. 5c-d)**. These results indicate that TMSC-Scr has therapeutic effects on preventing and rescuing RGC from degeneration in primary open angle glaucoma model.

**Fig. 5.**
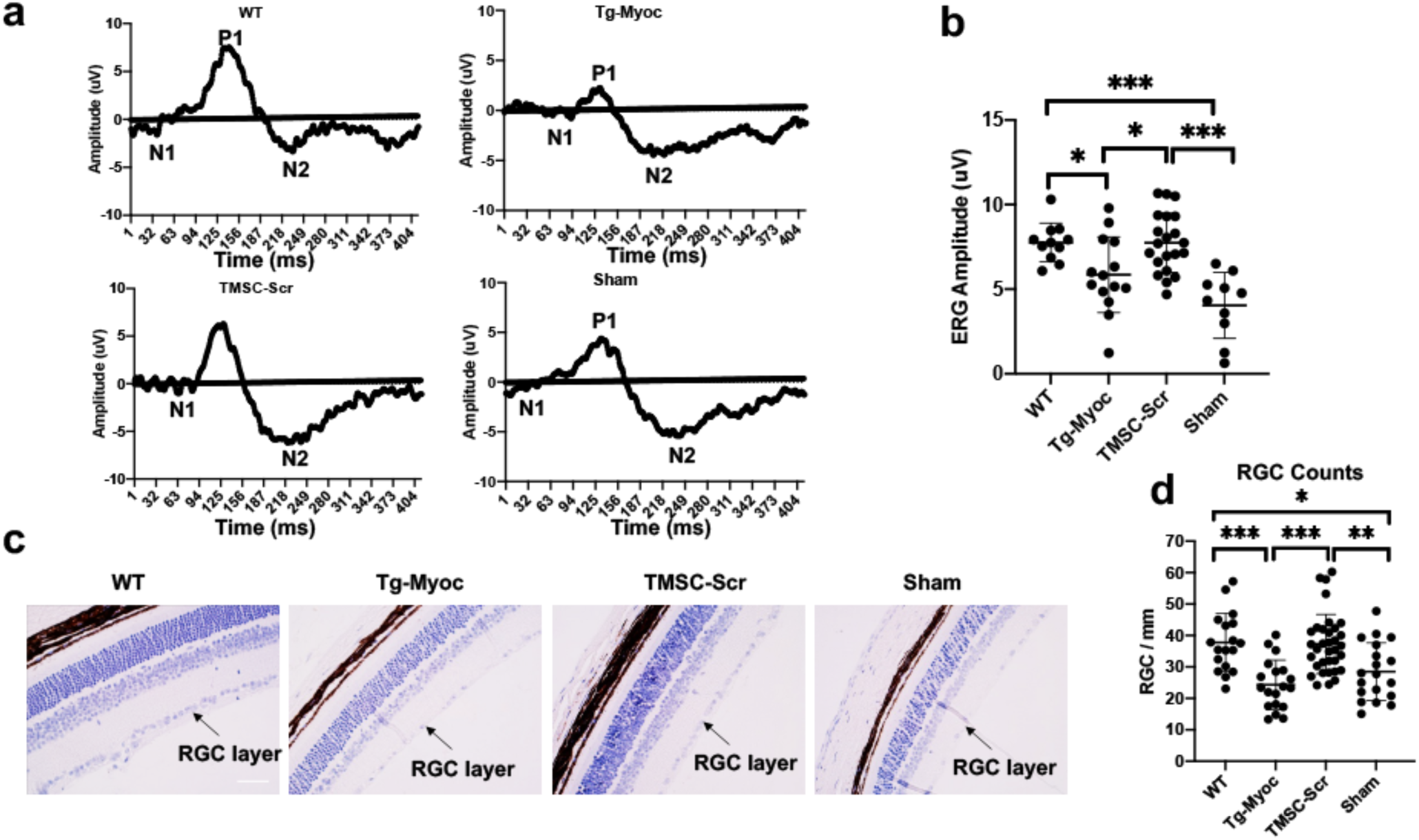
TMSC Secretome rescues RGC and retains function in Tg-MyocY437H mice. **a**. Pattern Electroretinogram (PERG) peaks showing amplitude of waves with P1 amplitude as an indicator of RGC function (n=11-21), **b**. Bar diagram showing average amplitude of PERG P1-wave peaks obtained for each group, **c**. DIC monocolor images showing RGC layer in Tg-MyocY437H mice, **d**. Bar diagram showing quantification of RGC number in Tg-MyocY437H mice retina, (n=3-4). Multiple dots in bar graphs represent combined results of biological and technical replicates, scale bar-100 µm. *P<0.05, **p<0.001, ***P<0.0001. Mean±SD, One-way ANOVA followed by Tukey posttest.

### TMSC secretome is enriched with proteins involved in ECM remodeling

We carried out the label-free proteomic identification and quantification of secretome proteins from two strains of human TMSC from different donors and compared them with the secretomes from two fibroblast strains. As observed by pathway enrichment and comparative analysis, TMSC-Scr showed upregulation of important proteins related to unfolded protein response (UPR), ECM organization proteins, and collagen catabolic process proteins (**Fig. 6a-c)**. STRING analysis of TMSC-Scr showed interaction patterns between proteins involved in unfolded protein response, cell matrix adhesion, and response to hypoxia detoxification **(Fig. 6d)**. Functional analysis identified 15 proteins in TMSC-Scr which were involved in promoting cell proliferation and maintenance of stemness in progenitor cells (NUDC, NAP1L1, NENF, MYDGF, etc.), while only 6 of these proteins could be identified in Fibro-Scr (**Supplementary Table 1**). These results show that the TMSC-Scr is enriched with the proteins important for TM cell survival, imparting protection against glaucoma.

**Fig. 6.**
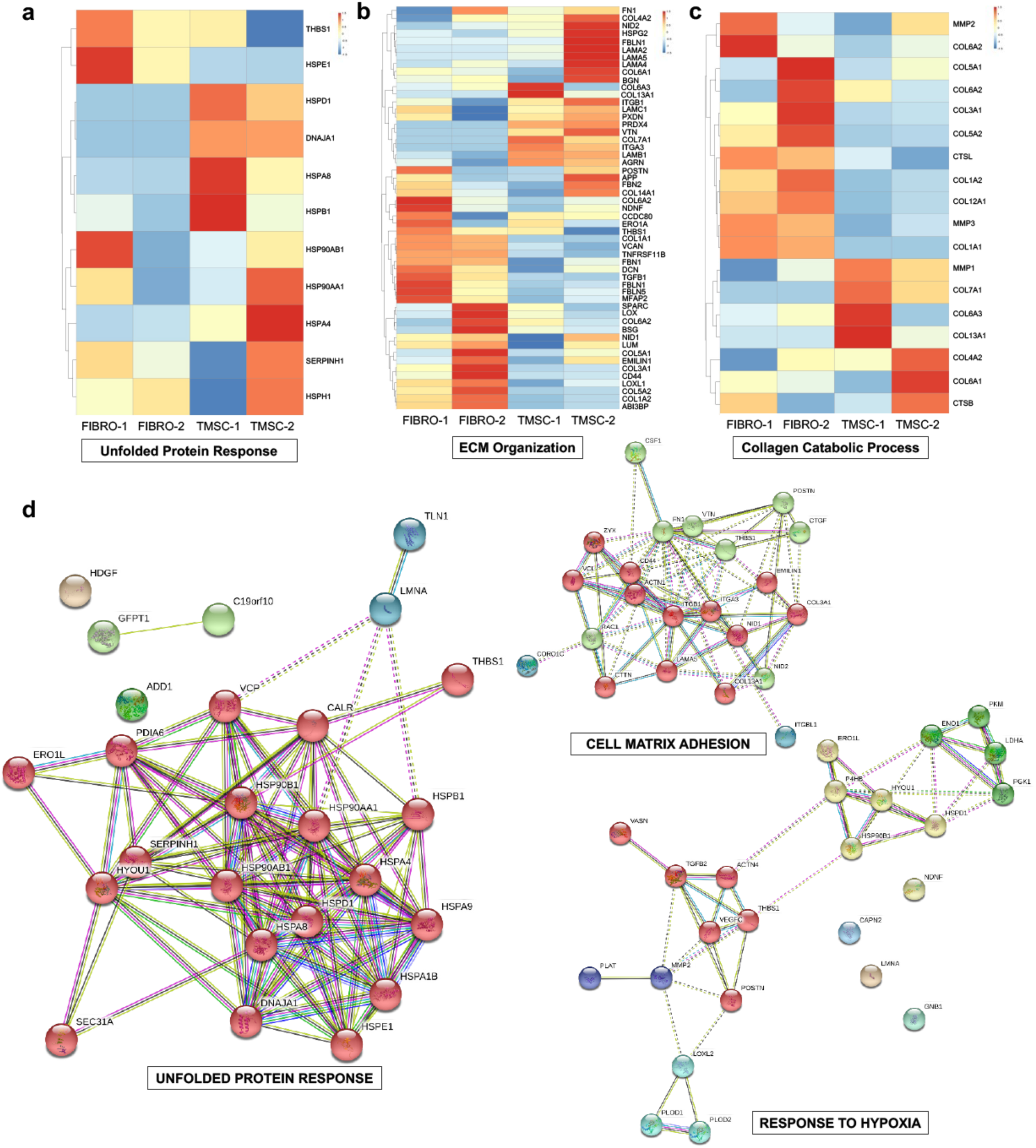
TMSC secretome contains different regenerative proteins evaluated by proteomics. **a-c**. Heatmaps and hierarchical clustering analysis showing differences in secretome proteins related to response to unfolded protein response, ECM organization, and collagen catabolic process between TMSC-Scr and fibroblast-Scr, **d**. Interactome analysis showing interaction between main proteins presented in TMSC-Scr, unfolded protein response, cell matrix adhesion, and response to hypoxia as analyzed by String v11, (n=2 each).

## Discussion

In this study, we explored the therapeutic potential of secretome from human TMSC and discovered potential mechanisms. TMSC-Scr was able to prevent as well as reverse Dex-induced TM cell changes in culture. By minimal invasive periocular injection, TMSC-Scr reduced the IOP, increased the TM cellularity, remodeled the ECM of the TM, activated the endogenous stem cells, promoted cell proliferation, and prevented and reversed RGC loss as well as preserved the RGC function in both steroid-induced and genetic myocilin mutant mouse models. The rejuvenating and therapeutic effects of TMSC-Scr on glaucoma are associated with the activation of COX2-PGE2 signaling. Our study indicates the feasibility of stem cell-free therapies for glaucoma in restoring both the TM function and RGC function for IOP reduction and vision preservation.

Ocular hypertension induced by steroids is a common side-effect among patients using steroid therapy. Myocilin mutations have been reported to be the most common form of genetic glaucoma^27^. We found that Dex treatment by periocular injection into C57BL/6J mice leads to increased fibrotic ECM proteins, such as fibronectin as well as myocilin in the TM resulting in protein misfolding, ER stress, and IOP elevation. Transgenic mouse model with myocilin Y437H mutation is characterized with increased IOP, persistent TM ER stress, RGC loss and axonal degeneration, which resembles POAG in patients ^20^. We found all these characteristics of the Tg-MyocY437H mice in this study. These two mouse models represent typical steroid-induced and genetic glaucoma. TM cell loss is associated with elevated IOP in glaucoma and reducing IOP is the only effective treatment so far. Our recent study reported the rescuing of glaucomatous conditions via improvement of TM homeostasis and prevention of RGC death in Tg-MyocY437H mice^10^. We investigated that TMSC-Scr treatment led to reduced IOP in both mouse models of glaucoma, which started as early as the following week of secretome periocular injection and persisted through experiment termination two to three weeks after secretome withdrawal.

CHI3L1 and AQP1 are involved in TM cell function of remodeling ECM, maintaining TM homeostasis, and modulating aqueous outflow^8,22,30^. After Dex-Ac treatment, TM cells lose these proteins and become stiffer^45^. Increased CHI3L1 and AQP1 and reduced glaucoma associated genes *MYOCILIN* and *ANGPTL7* after TMSC-Scr treatment in Dex-treated TM cells reflect the rejuvenating ability of TMSC-Scr on TM cells. Postoperative fibrosis is a major complication in glaucoma, and fibronectin is the main fibrotic protein^46^. TMSC-Scr treatment in Dex-treated cells provides an evidence for TMSC-Scr mediated fibrosis reduction.

Additionally, secretome treatment increased the number of TM cells reflecting its ability to regenerate TM tissue *in vivo*. Collagen forms major connective tissue protein in ECM and its degradation by matrix metalloproteinases (MMPs) is crucial for remodeling and repairing of tissue^47^. TMSC-Scr treatment dramatically downregulated the fibrotic ECM components fibronectin and collagen IV, which might help TM cell survival. The TMSC-Scr proteins involved in ECM organization and collagen catabolic process could lead to enhanced ECM turnover and lower IOP. ER stress of the TM cells is involved in both glaucoma models. To counteract protein misfolding, cells activate the UPR pathway. ER chaperones GRP78 and CHOP are activated by UPR resulting into restoration of ER homeostasis by proteasome mediated induction of ER associated degradation^48^. In our study, the presence of UPR in TMSC-Scr might further induce the cellular responsiveness of TM cells to reduce ER stress.

Prostaglandin analogs are commonly used for glaucoma treatment to reduce IOP. COX2, a rate-limiting enzyme is known to convert arachidonic acid to PGH2, which is isomerized to PGE2 by PGE2 synthase^36^. COX2 modulates PGE2 synthesis in response to growth factors, inflammatory cytokines, and other physiological demands. Under normal conditions, COX2 is restricted mostly in kidney^49^. COX2 expression can be increased substantially in other tissues in response to proinflammatory cytokines or sheer stress^50^. COX2 expression is completely lost in the non-pigmented secretory epithelium of ciliary body and aqueous humor of end-stage POAG human eyes^51^. Glucocorticoids are well known for inhibiting COX2 activity^52^. Human TM cells can secrete PGE2 which was inhibited significantly after a moderate Dex treatment. Mitochondrial TMEM177 has also been reported to associate with COX20 to stabilize/increase the biogenesis of COX2^39^. Hence, increased TMEM177 after TMSC-Scr treatment might be responsible for increased COX2 stabilization and biogenesis, which further indicates its therapeutic potential for glaucoma. Further evaluation of increased TMEM177/COX2 expression in Tg-MyocY437H mice after TMSC transplantation confirmed that both stem cell-based and cell-free therapy involves upregulation of COX2 to impart a therapeutic benefit. Abrogation of COX2 by treated with inhibitors resulting in the loss of protective effect of TMSC-Scr on TM cells provides further evidence for important role of COX2-PGE2 signaling in secretome mediated TM protection. Further investigations in this direction might help to explore the effect of COX2 inhibition on the levels of MMPs and thus remodeling of glaucomatous ECM.

Mobilization and maintenance of endogenous stem cells is very crucial for inducing tissue regeneration^53^. Endogenous stem cell regeneration involves a complex interplay of cues in terms of growth factors, modulation in stem cell niche, and chemokines inducing differentiation, proliferation, and migration of these cells. Glaucoma and ER stress have been reported to decrease endogenous stem cells and increase apoptosis^11^. PGE2 has been reported to increase self-renewal and proliferation of stem cells^40^. Recently Pala et al. discovered the interesting role of PGE2 in restoring muscle function in aging induced animals in addition to augmenting stem cell function^54,55^. PGE2 has been also reported to increase hematopoietic^56^ and bone homeostasis^57^ and aid in liver regeneration^58^. Our study provides an additional supporting evidence that PGE2 also induces a regenerative effect in the eye disorders like glaucoma by increasing stem cell activation and function. Increased PGE2 secretion in aqueous humor of both mouse models after TMSC-Scr treatment might be crucial for maintaining the ABCB5^+^ and OCT4^+^ endogenous stem cells and reversing glaucomatous changes. The stem cell proliferation and renewal are further enhanced by TMSC-Scr by the presence of crucial proteins involved in promoting cell proliferation and stemness in progenitor cells. Zhu et al reported that intracameral injection of iPSC-derived TM cells in Tg-MyocY437H mice increased TM cellularity and reduced IO mainly by activation of endogenous TM cells^14,59^ Although the endogenous TM cells and TMSC in Tg-MyocY437H mice should carry the transgenic abnormal gene, whether stem cells and stem cell secretome improve their function needs further investigation.

Hypoxia can lead to RGC death by inducing a number of degenerative changes. The increased secretion of response to hypoxia proteins in TMSC-Scr might be responsible for RGC survival/rescue in hypoxic conditions. Uncovering the key proteins behind neuroprotective effect of TMSC-Scr on RGC survival and function can be an interesting avenue for future investigations. Furthermore, therapeutic effects of TMSC-Scr given through the periocular route emphasize that secretome proteins can cross the corneal barrier and can be a good approach for development of glaucoma eye drops.

Stem cell secretome therapy can be implicated soon for clinical trials after studying confirmation in animals more relevant to humans like non-human primates. The key secretome proteins that have been identified in our current study might potentially lead to designing small molecule-based therapeutics for glaucoma in the future.

## Materials and Methods

### Study Design

The primary objective of this research study was to evaluate the effect of human TMSC secretome in steroid induced and genetic mouse models of glaucoma. All the experiments conducted on the animals were approved by the University of Pittsburgh Institutional Animal Care and Use Committee and complied with the ARVO Statement for the Use of Animals in Ophthalmic and Vision Research. Four-months old wildtype (WT) C57BL/6J mice were purchased from Jackson Laboratory as wildtype control (WT). Adult C57BL/6J mice were periocularly injected with 20 µl of dexamethasone acetate (Dex-Ac) (DE122, Spectrum Chemicals) at 10 mg/ml once a week following the published protocol^19^. The genetic POAG model, transgenic mice with myocilin Y437H mutation (Tg-MyocY437H) were kindly gifted from Gulab Zode (North Texas Eye Research Institute, Texas)^20^ and bred with C57BL/6J WT mice and genotyped to confirm the myocilin mutation. Littermates were used as controls. Primary human TMSC, TM cells and corneal fibroblasts were derived from deidentified corneas unsuitable for corneal transplantation obtained from the Center for Organ Recovery and Education (CORE, Pittsburgh, PA) or from the corneal rims after corneal transplantation. Proper informed consent for using the donated corneas for research purpose was obtained from all the donors by the CORE. The study was performed in accordance with the proper guidelines and regulations set by the University of Pittsburgh and under an IRB-exempt protocol and IBC protocol approved by the University of Pittsburgh. Blinded methods were used for study outcome and designs. In particular, the histological evaluations of cell immunofluorescence for different antibodies and TM and RGC cell counts were performed by two independent researchers blindly. IOP were measured by three independent researchers. qPCR experiments were performed by another researcher in a blinded manner. We estimated the sample size based on the variability of different assays and potential outliers. Sample size for Dex-Ac and Tg-MyocY437H mice cohorts was estimated by power analysis before initiation of the study. Number of samples (n) used for each experiment is specified in the captions for each figure.

### Primary cell culture and characterization

TMSC were cultured in Opti-MEM (Invitrogen) with supplements including 5% fetal bovine serum (FBS), 0.08% chondroitin sulfate, 100 μg/ml bovine pituitary extract (ThermoFisher), 20 μg/ml ascorbic acid, 10 ng/ml epidermal growth factor, 200 μg/ml calcium chloride (Sigma-Aldrich), 50 mg/ml gentamicin, 100 mg/ml streptomycin, and 100 IU/ml penicillin (ThermoFisher) as published previously^8,21^. Stem cell characterization was performed similar to our published method^22^. TM cells were cultivated in DMEM: HAM’s F12 (1:1) medium with 10% FBS and confirmed by responsiveness to 100 nM Dex^25,26^. TMSC were used between passages 4-7. Secretome treatment was given 1) in cell culture together with Dex (referred as “parallel TMSC-Scr”) for five days (for preventive effect); 2) TM cells were treated with Dex for five days and then secretome was supplemented in cell culture in the presence of Dex for another five days (referred as “post TMSC-Scr”) (for reverse effect). Human corneal fibroblasts were collagenase digested and cultured in DMEM/F12 with 10% FBS and used between passage 4-7 as described previously ^24,60,61^. For COX2 inhibition assay, dex and secretome treated primary TM cells were treated additionally with Nimesulide (200µM) and Indomethacin (100µM) (Sigma Aldrich) for ten days. Media was changed every third day. For immunofluorescent staining, cells were grown on coverslips in triplicates. Three different TM cells were treated with secretome from two different TMSCs. All antibodies used are listed in **Supplementary Table 2. Flow cytometry**

For stem cell characterization, TMSC were blocked in 1% bovine serum albumin (BSA) for one hour and incubated with fluorochrome conjugated antibodies for 30-min on ice in dark. 5′104 cells were acquired per tube on BD FACS Aria (BD Biosciences). Compensations were adjusted using proper controls and isotype controls were used to rule out spectral bleeding. Cell apoptosis was assessed by Annexin V/7-AAD staining. Data was analyzed using the FlowJo_V10 software (FlowJo).

### Calcein AM/Hoechst staining

Post secretome harvesting (∼90% confluence), TMSC and corneal fibroblasts were stained for 15 minutes in dark with the viability dyes Hoechst 33342 (1:2000) and Calcein AM (1:1000) (Invitrogen). Live cells were captured at excitation filters of wavelength 361nm and 565nm respectively employing TE 200-E (Nikon).

### MTT assay

To assess the cell proliferation post secretome harvesting, 5×103 TM cells were cultured per well in 96-well plates in optimum culture condition as described above. These cells were then incubated with secretome from three different TMSC lines for 48h. MTT reagent (Sigma-Aldrich) was used to assess the formation of formazan crystals at end point. The optical density was measured via ELISA reader (Tecan) using 570 nm wavelength and considering 600 nm as reference to eliminate any background noise. Cells grown in optimum culture media without secretome were taken as control.

### Eye section processing and staining and cell counting

Whole mouse eyes were enucleated into freshly prepared 1% PFA in PBS and fixed for at least 48h at 4°C. For plastic sectioning, eyes were embedded in glycol methacrylate A resin (JB-4, Polysciences) according to manufacturer’s instructions. Specifically, the eyes were placed in 70% ethanol and then dehydrated in increasing concentrations of ethanol to 100%. Fully dehydrated samples were infiltrated with resin monomer and its catalyst overnight. The next day, accelerant was added and the samples embedded. Then 3-µm sections were made on a rotary microtome (RM2235, Leica). Slides were immediately stained for 1 minute in Mayer’s hematoxylin solution (Electron Microscopy Sciences). Sections were photographed in DIC color mode using 40X oil objective (Olympus). For RGC counting, retina was imaged at nasal and temporal sides and three images were captured per eye. RGC counting was done in 3-4 eyes per group in both Dex-Ac and Tg-MyocY437H models using automated cell counter mode in ImageJ (NIH). Total counted RGC in a defined length of retina were normalized to calculate cells/µm for each group.

For cryosections, fixed whole mouse eyes were embedded sagittally in OCT compound (Tissue Plus, FisherScientific) and flash frozen in liquid nitrogen chilled isopentane. 10-µm sections through the center of the eye were made on a motorized Cryostat (CM3050 S, Leica).

### Immunofluorescent staining

Cells were cultured on glass coverslips and fixed with 4% paraformaldehyde (PFA) (Electron Microscopy Sciences). Fixed cells were permeabilized with 0.1%Triton X-100 and blocked with 1% BSA and stained with corresponding primary antibodies overnight at 4°C. After washing with PBS, cells were stained with specific secondary antibodies conjugated with fluorochromes FITC, PE, and APC. Nuclei were stained with DAPI. Samples were acquired under a confocal laser scanning microscope (Olympus).

### Quantitative Real-Time PCR (qPCR)

Cells were lysed with RLT buffer and total RNA was isolated using an RNA purification kit (RNeasy Mini Kit, Qiagen). cDNAs were transcribed using High-Capacity cDNA Reverse Transcription Kit (Applied Biosystems). qPCR was performed by direct dye binding (SYBR Green, Applied Biosystems). Primers were designed using Primer3 and blasted on the NIH website (https://www.ncbi.nlm.nih.gov/tools/primer-blast/) to confirm the specificity. The primer sequences were: *MYOCILIN* (forward: AAGCCCACCTACCCCTACAC; reverse: TCCAGTGGCCTAGGCAGTAT), *ANGPTL7* (forward: GCACCAAGGACAAGGACAAT; reverse: GATGCCATCCAGGTGCTTAT). RNA content was normalized by 18S rRNA (Forward: CCCTGTAATTGGAATGAGTCCAC, Reverse: GCTGGAATTACCGCGGCT).

Relative mRNA abundance was calculated as the Ct for amplification of a gene-specific cDNA minus the average Ct for 18S expressed as a power of 2 (2^ΔΔCt^). Three individual gene-specific values thus calculated were averaged to obtain mean ± SD.

### Secretome preparation

Secretomes were harvested from both TMSC and corneal fibroblasts. 1×10^6^ cells were cultured per T75 flask in the log phase and incubated with basal media without serum and growth factors at 60-70% confluence for 48h. Then, the culture supernatant was centrifuged at 3000rpm for 5 minutes to remove any cell debris and concentrated using 3kDa centricon devices (Amicon) to 25X and immediately stored at -80°C until use.

### Secretome dosing and treatment

For cell culture, 1x secretome was mixed with basal DMEM/F12 (1:1) in 100 nM Dex (Sigma-Aldrich) treated TM cells. For animal experiments, 20µl of 25x concentrated secretome was periocularly injected to each mouse eye. In Dex-Ac model, both TMSC and fibroblast secretomes were injected at week 3, continued once a week till week-6 and animals were sacrificed at week-8. In Tg-MyocY437H model, TMSC-Scr was injected at week 0 when mice were 4 months old, continued once a week till week-7, and animals were sacrificed at week-10. IOP was measured once a week.

### Periocular secretome injection and IOP measurement

Two TMSC strains from two different donors at passages 4-7 were used for secretome isolation and mouse injection. Fibro-Scr was used as a control in the Dex-Ac induced model. For the Dex-Ac model, mice were divided in four groups: Dex-Ac treated mice (n=16), vehicle injected with vehicle solution used for solubilizing Dex-Ac (n=16), TMSC-Scr group (n=16), and Fibro secretome group (n=16). For genetic model, mice were divided into five groups: littermate group (WT, n=16), Tg-MyocY437H mice (Tg, n=16), Tg-MyocY437H mice with periocular injection of the basal medium (Tg-Sham, n=16), and Tg mice with periocular injection of TMSC-Scr (Tg-TMSC, n=16 each). For TMSC injections, Tg-MyocY437H mice were divided into four groups as described above (n=3-4 per group) and injected with 5×10^4^ DiO-green labelled TMSC intracamerally. In brief, mice were anesthetized with ketamine-xylazine^8,9,13^. 20μl of 25x concentrated secretome or basal medium was injected periocularly using a 33-gauge needle connected to a Hamilton syringe. An I-care tonometer was used to measure mouse IOP (TonoLab; Colonial Medical Supply). All IOP measurements were performed between 1 PM and 3 PM. IOP measurement before injection served as baseline and measurements were conducted once a week up to 8-10 weeks.

### Pattern Electroretinography (PERG)

PERG was performed on the Celeris (Diagnosys) to evaluate the RGC function. Mice (n=9-15 for each group) were dark adapted overnight and anesthetized with intraperitoneal injection of Ketamine and Xylazine. Mouse pupil was dilated with 0.5% Tropicamide and 2.5% Phenylephrine eye drops. A circular electrode centered on the cornea was placed in a plane perpendicular to the visual axis after applying GenTeal lubricant to avoid corneal dryness and prevent cataract formation. Pattern stimuli consisted of horizontal bars of variable spatial frequencies and contrast that alternate at different temporal frequency. The parameters for PERG amplitude were spatial frequency 0.155 cycles/degree, temporal frequency 2.1 reversals/sec, contrast 100% and substantial averaging (600-1800 sweeps). The Amplitude of P1 wave was used to analyze the function of RGC.

### Immunoblotting

Mouse limbus tissue and aqueous humor were lysed using RIPA buffer (SantaCruz Biotechnology). Limbus tissue was cut and sonicated to fine pieces. For secretory Myoc, aqueous humor was extracted from the eyes. Protein concentration was measured using the BCA Protein Assay Kit (Pierce Biotechnology). Protein samples were loaded for each group in each well and run on 8-16% sodium dodecyl sulfate–polyacrylamide gel (ThermoFisher) for electrophoresis and then transferred to the PVDF membrane. After blocking with blocking buffer, membranes were incubated with primary antibodies overnight at 4°C. Corresponding secondary antibodies (IRDye 680LT and IRDye 800CW, LI-COR Biosciences) were incubated after three washes of 0.1% Tween 20 in Tris-buffered saline. Detection and capture of fluorescent signal were performed using an infrared imager (Odyssey; LI-COR Biosciences). ImageJ was used for the quantification and analysis of protein expression with β-actin as internal control.

### Enzyme linked Immunosorbent Assay (ELISA)

Secreted PGE2 was measured in aqueous humor samples using PGE2 ELISA (Enzo life Sciences) and procedure was performed according to manufacturer’s instructions. Aqueous humor was used at a dilution of 2:100 for the assay. Optical density was measured at 405nm using 570nm and 600nm as normalizing wavelengths using ELISA reader. Optical density measurements were extrapolated to measure PGE2 concentration (pg/ml) reference to standards.

### Multidimensional protein identification technology (MudPIT) analysis

TCA-precipitated proteins were urea-denatured, reduced, alkylated and digested with endoproteinase Lys-C (Roche) followed by modified trypsin (Promega)^62,63^. Peptide mixtures were loaded onto 250 μm fused silica microcapillary columns packed with strong cation exchange resin (Luna, Phenomenex) and 5-μm C18 reverse phase (Aqua, Phenomenex), and then connected to a 100 μm fused silica microcapillary column packed with 5-μm C18 reverse phase (Aqua, Phenomenex)^62^. Loaded microcapillary columns were placed in-line with a Quaternary Agilent 1100 series HPLC pump and a LTQ orbitrap Elite mass spectrometer equipped with a nano-LC electrospray ionization source (ThermoScientific). Fully automated 10-step MudPIT runs were carried out on the electrosprayed peptides. Tandem mass (MS/MS) spectra were interpreted using ProluCID v. 1.3.3 against a database consisting of 79096 nonredundant human proteins (NCBI, 2016-06-10 release), 193 usual contaminants. To estimate false discovery rates (FDRs), the amino acid sequence of each non-redundant protein entry was randomized to generate a virtual library. This resulted in a total library of 158578 non-redundant sequences against which the spectra were matched. All cysteines were considered as fully carboxamidomethylated (+57.0215 Da statically added), while methionine oxidation was searched as a differential modification (+15.9949 Da). DTASelect v 1.9^64^ and swallow v. 0.0.1 (https://github.com/tzw-wen/kite), an in-house developed software, were used to filter ProLuCID search results at given FDRs at the spectrum, peptide, and protein levels. Here all controlled FDRs were less than 5%. All 4 data sets were contrasted against their merged data set, respectively, using Contrast v 1.9 and in house developed sandmartin v 0.0.1. Our in-house developed software, NSAF7 v 0.0.1, was used to generate spectral count-based label free quantitation results^65^.

### GO enrichment analysis

DAVID software, a free online tool (version DAVID 6.8; http://david.ncifcrf.gov/), and GOstats (version 2.48.0) were used for the functional enrichment analysis of genes whose corresponding proteins had positive distributed normalized spectral abundance factor (dNSAF) values in both replicates of a specific cell type. Biological process (BP); cellular component (CC); and molecular function (MF) were three different categories according to which the GO terms classification was performed. The top 10 most significantly enriched GO categories for a given cell type were compared between fibroblasts and TMSC cells.

### Hierarchical clustering analysis and Heatmap

The hierarchical clustering analysis and heatmap plotting was performed using the R package pheatmap (version 1.0.12). Comparative expression of the dNSAF protein values between TMSC and fibroblast cells were performed using heatmaps. For each row of the heatmap, the name of the gene that encodes the protein was used as the heatmap row name. Similar elements were classified in groups in a binary tree using hierarchical clustering.

### Statistical Analysis

Results were expressed as mean ± standard deviation (SD). The statistical differences were analyzed by Two-way ANOVA (for the IOP data) or one-way ANOVA (all others, except protein expression in human glaucomatous tissue in Figure S5, unpaired t-test), followed by Tukey posttest using PRISM. p< 0.05 was considered statistically significant.

## Supporting information

Supplemental Tables and Figures in one pdf file

## Acknowledgements

We thank all donors and their families for donating corneas and the clinicians at UPMC Eye Center Drs. Deepinder Dhaliwal, Vishal Jhanji, Alex Mammen, and the fellows and residents who supplied the corneal rims, and the Center for Organ Recovery & Education (CORE) in Pittsburgh who supplied the corneas. We thank Gulab Zode for providing Tg-MyocY437H mice. We thank Nancy Zurowski for Flow Cytometry. Fig. 1a, Fig. 3a, and graphical abstract were created with BioRender.com.

## Author contributions

Conceptualization and designing of work, A.K., Y.D.; Acquisition, analysis, and interpretation of the data, A.K., Y.D., T.X., A.P.; Investigation, A.K., X.S., M.Z., W.C., E.Y., A.P., L.L., A.P., Y.Z., L.F., M.W., Ak.K., Y.L., Y.X., K.L., K.D., Y.D.; Writing-Original Draft, A.K., Y.D.; Writing-Review & Editing, A.K, K.L., K.D., Y.C., J.S.S., T.X., Y.D.; Funding Acquisition, Y.D.; Resources, Y.D. and T.X.; Supervision, Y.D. All authors have read and approved the manuscript.

## Data and materials availability

All data associated with this study are provided in the paper or the Supplementary Materials. The primary human TMSC, TM, and fibroblast cell lines generated in this study are available from the Lead Contact with a completed Material Transfer Agreement. All label free proteomic quantification data had been submitted to Massive. After publication (i.e. the dataset is made public), the permanent URL to the dataset will be published and accessible to everyone.

## Supplementary Materials

Supplementary Fig. 1. TMSC characterization and evaluation of cell viability in corneal fibroblasts and TMSC post secretome harvesting.

Supplementary Fig. 2. TMSC secretome induces regeneration in Dex-induced TM cells

Supplementary Fig. 3. TMSC secretome modulates myocilin and ECM in Tg-MyocY437H.

Supplementary Fig. 4. Secretome treatment leads to increased recruitment of endogenous TM stem cells.

Supplementary Fig. 5. TMSCs and COX2 are reduced in human glaucoma donor TM tissue

Supplementary Table 1. List of the proteins directly involved in promoting cell proliferation and maintenance of stemness in progenitor cells, identified in secretome of TMSC and fibroblasts.

Supplementary Table 2. List of all the antibodies used in the study.

Data file 1: Original Immunoblotting gels (provided as separate pdf file) with clear indication of which bands were used in the figures in the form of arrows.

