## Supplemental Tables and Figures in one pdf file for "Stem cell-free therapy for glaucoma to preserve vision"

**This file includes:**

**Supplementary Fig. 1.** TMSC characterization and evaluation of cell viability in corneal fibroblasts and TMSC post secretome harvesting.

**Supplementary Fig. 2.** TMSC secretome induces regeneration in Dex-induced TM cells

**Supplementary Fig. 3.** TMSC secretome modulates myocilin and ECM in Tg-MyocY437H.

**Supplementary Fig. 4.** Secretome treatment leads to increased recruitment of endogenous TM stem cells.

**Supplementary Fig. 5.** TMSCs and COX2 are reduced in human glaucoma donor TM tissue.

**Supplementary Table 1.** List of the proteins directly involved in promoting cell proliferation and maintenance. of stemness in progenitor cells, identified in secretome of TMSC and fibroblasts.

**Supplementary Table 2.** List of all the antibodies used in the study.

**Supplementary Figures**

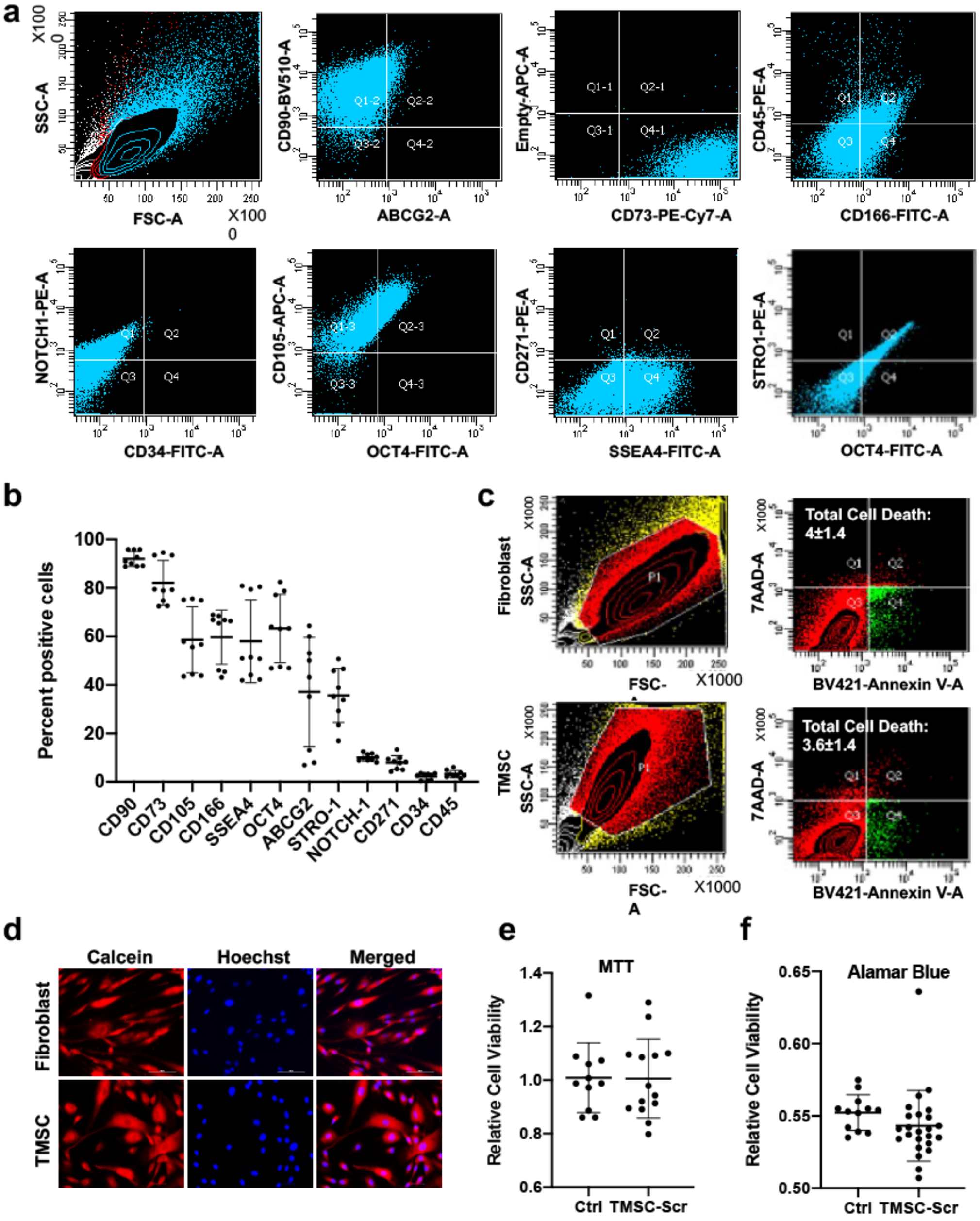


**Supplementary Fig. 1. TMSC characterization and evaluation of cell viability in corneal fibroblasts and TMSC post secretome harvesting. A-b.** Dot plot and bar diagram showing percent positivity for different stem cell markers of TMSC, **c.** Annexin V/7-AAD flow cytometry analysis for cell viability assessment in corneal fibroblasts and TMSC post secretome harvesting, after incubation in serum free media. Gate was set on Unstained control cells for normalizing the background fluorescence. 100,000 events per tube were acquired. Results are representative of Mean±SD of three independent experiments (n=3), **d.** Live cell fluorescent microscopy for cell viability in corneal fibroblasts and TMSC post secretome harvesting. Calcein AM and Hoechst 33342 were used to stain viable cells. **e.** MTT assay, and **f.** Alamar blue assay showing maintenance of cell viability after secretome incubation for 48 hours in TM cells, an indication of non-toxicity of secretome in the TM cells, (n=4). Multiple dots in bar graphs represent combined results of biological and technical replicates. Student's t-test, scale bar-100 µm.

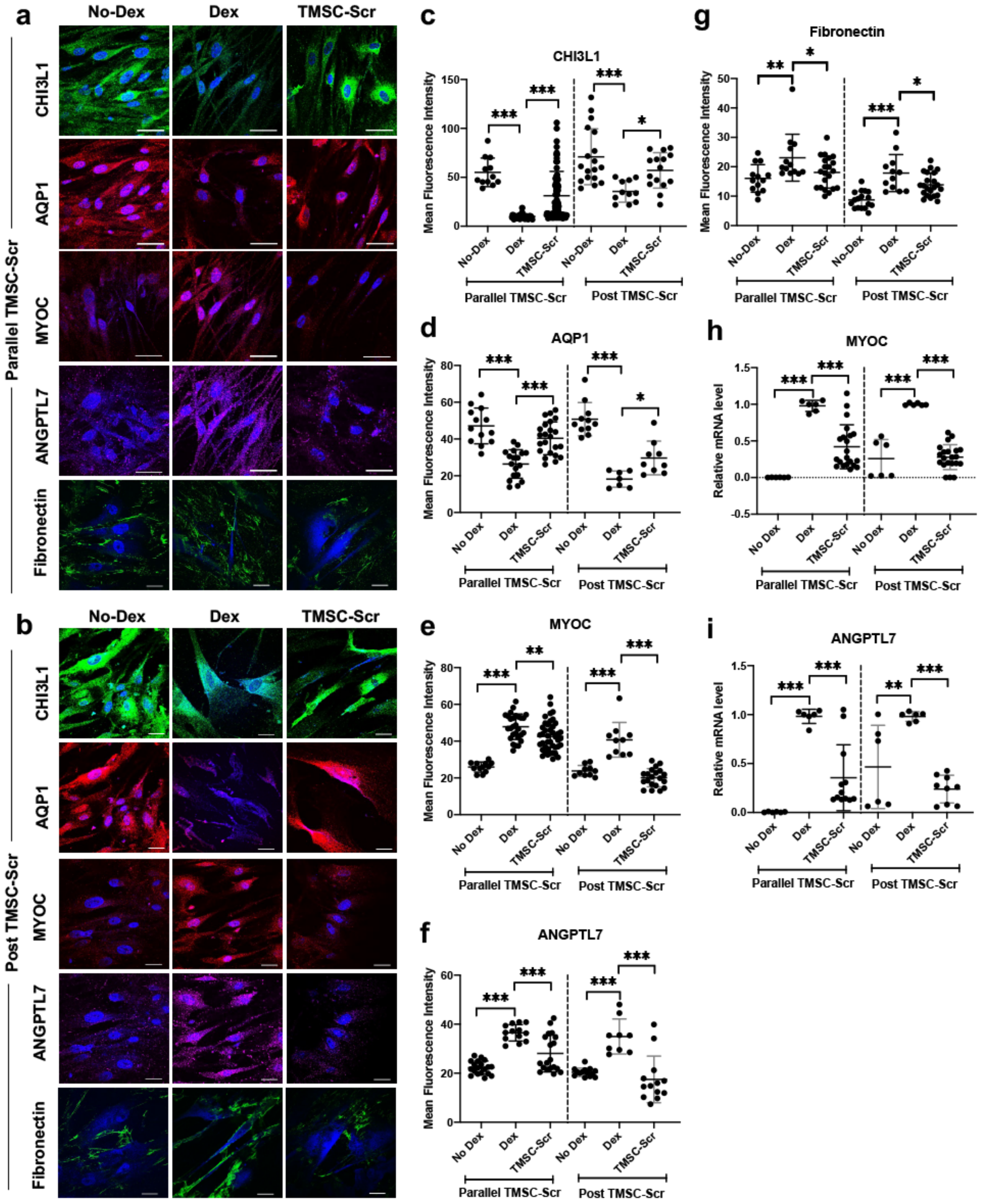


**Supplementary Fig. 2. TMSC secretome induces regeneration in Dex-induced TM cells. a-b.** Immunofluorescent pictures showing protein expression of CHI3L1, AQP1, Myoc, ANGPTL7 and fibronectin in TMSC-Scr treated cells in parallel and post to Dex induction, **C-G.** Bar diagrams showing quantification of mean fluorescent intensity of the staining in **a-b**, **h-i.** Bar diagrams showing relative mRNA expression of glaucoma associated genes *MYOC* and *ANGPTL7* comparing TMSC-Scr treated and untreated cells. Experiments were repeated with at least three primary TM cell strains with secretome used from four different TMSC strains. Multiple dots in bar graphs represent combined results of biological and technical replicates. Scale bar-50 µm (panel **a** except fibronectin), scale bar-30 µm (panel **a**-fibronectin and panel **b**), *p<0.05, **p<0.001, ***p<0.0001. Mean±SD, One-way ANOVA followed by Tukey posttest.

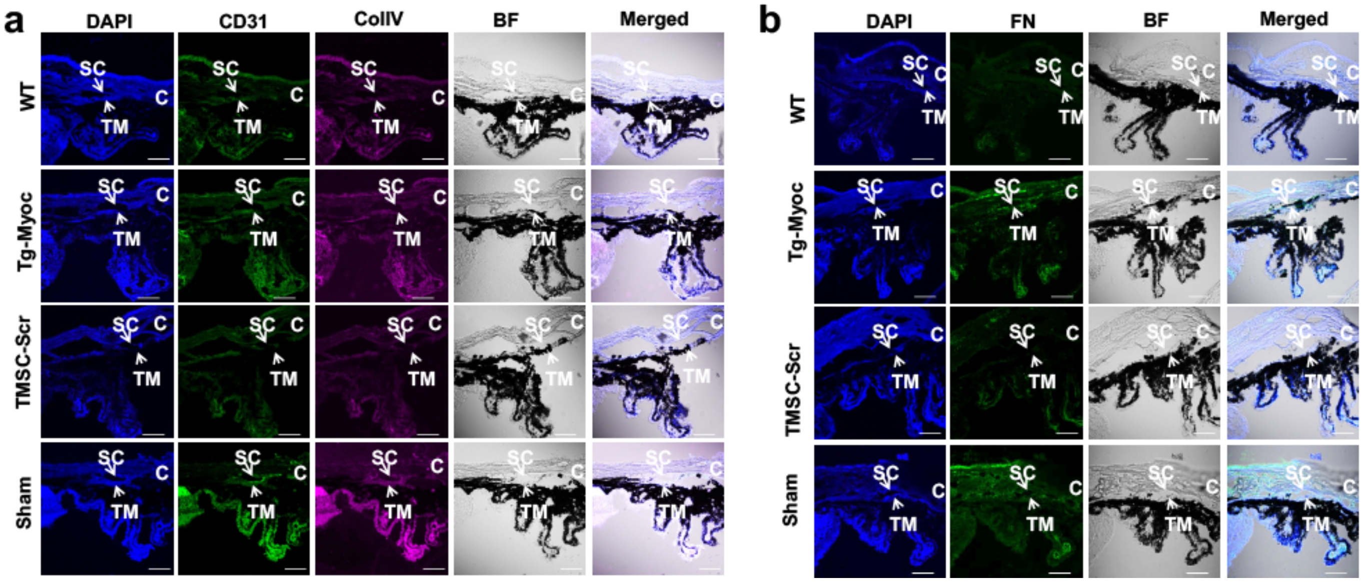


**Supplementary Fig. 3. TMSC secretome modulates myocilin and ECM in Tg-MyocY437H. a.** Immunofluorescent images showing protein expression of CD31 and ColV in Tg-MyocY437H mice, **b.** immunofluorescent images showing expression patterns of FN in Tg-MyocY437H mice, SC-Schlemm's canal, TM-Trabecular Meshwork, scale bar-30 µm.

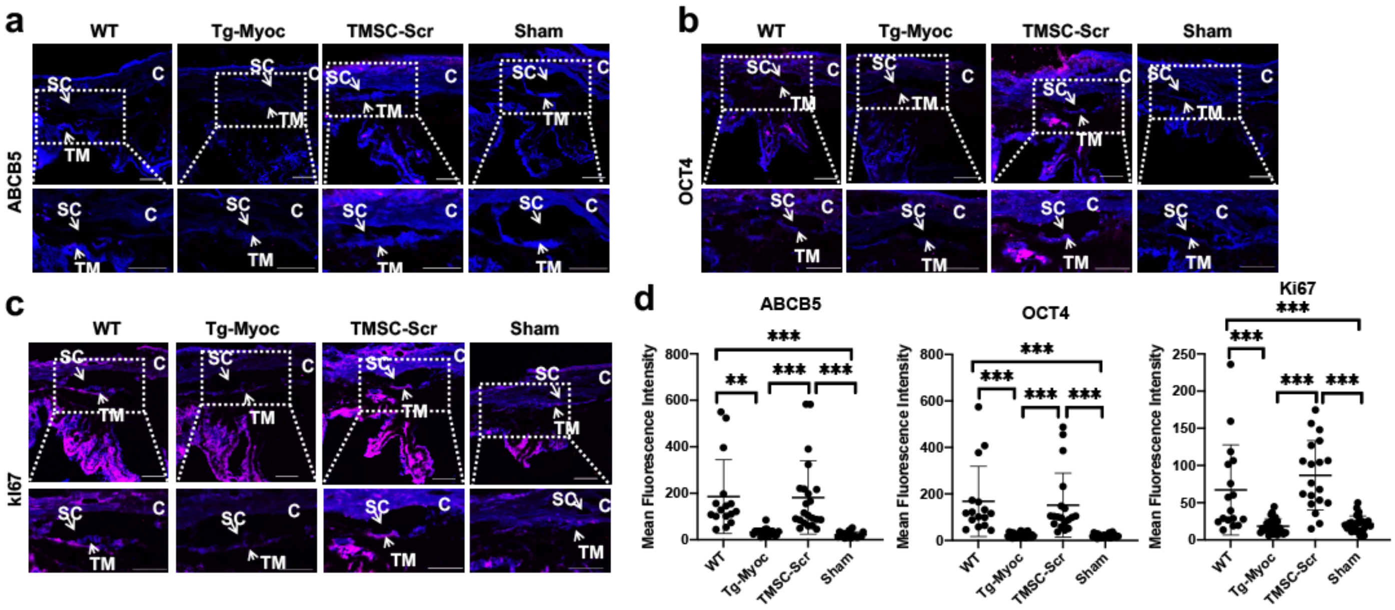


**Supplementary Fig. 4. Secretome treatment leads to increased recruitment of endogenous TM stem cells. a-d.** Immunofluorescent images and bar graphs showing protein expression and quantification of ABCB5, OCT4 and Ki67, scale bar- 50 µm. n=3-4 eyes per group, dots represent MFI quantified in multiple sections (4-16 limbus sections) per eye. *$p<0.05$, **$p<0.001$, ***$p<0.0001$. Mean±SD, One-way ANOVA followed by Tukey posttest. SC-Schlemm's canal, TM-Trabecular Meshwork, C-Cornea.

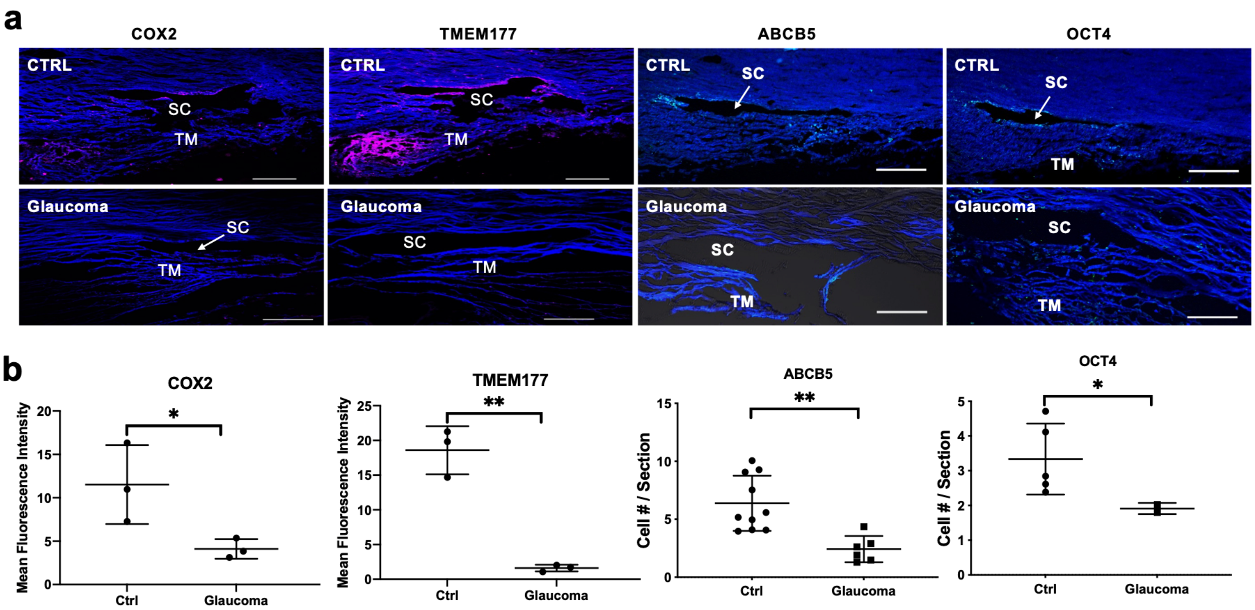


**Supplementary Fig. 5. TMSCs and COX2 are reduced in human glaucoma donor TM tissue. a.** A pair of the glaucomatous corneal tissues and 2 age-matched normal controls were cryosectioned and stained with ABCB5, OCT4, COX2, and TMEM177. Mean fluorescence intensity was quantified to compare the insert and filter region **b.** Dots represent section numbers of two tissues in each condition. Scale bars, 100 µm. *P<0.05, **p<0.001, ***. Mean±SD, unpaired t-test. SC-Schlemm's canal, TM-Trabecular Meshwork, C-Cornea.

**Supplementary Table 1.** List of the proteins directly involved in promoting cell proliferation and maintenance of stemness in progenitor cells, identified in secretome of TMSC and fibroblasts.

| Sr. No. | Accession No. | Name | TMSC | Fibro | Function |
|---|---|---|---|---|---|
| 1 | NP_009016.1 | FSTL1 | + | + | Modulates cell proliferation and helps in maintenance of stemness |
| 2 | NP_001054.1 | TF | + | - | Stimulates cell proliferation |
| 3 | NP_004416.2 | ECM1 | + | + | Increases cell proliferation and maintains stemness by stabilizing β-catenin |
| 4 | NP_006608.1 | NES | + | + | Required for mitogen stimulated proliferation and involved in self renewal of neural progenitor cells = |
| 5 | NP_002511.1 | NPM1 | + | + | Promotes proliferation by regulating ribosome biogenesis |
| 6 | NP_733821.1 | LMNA | + | + | Supports cell proliferation, knockdown induces cell senescence |
| 7 | NP_757351.1 | CSF1 | + | + | Induces proliferation in progenitor cells |
| 8 | NP_061980.1 | MYDGF or C19orf10 | + | - | Secreted by Bone marrow-cells to promote cardiac tissue repair and promotes cardiomyocyte proliferation and heart regeneration in neonate hearts. Stimulates endothelial cell proliferation through a MAPK1/3-, STAT3- pathway. Increase cardiomyocyte proliferation though PI3K/AKT-signaling pathway |
| 9 | NP_002473.2 | NASP | + | - | Required for cell proliferation |
| 10 | NP_006591.1 | NUDC | + | - | Highly expressed in human bone marrow myeloid and erythroid progenitors, induces cell proliferation; |
| 11 | NP_001294853.1 | NAP1L1 | + | - | Regulates embryonic neural progenitor cell proliferation |
| 12 | NP_005889.3 | CAPRIN1 | + | - | Increases cellular proliferation |
| 13 | NP_077719.2 | NOTCH2 | + | - | Mediates cell growth and prevents apoptosis |
| 14 | NP_006182.2 | PA2G4 | + | - | Maintains cell growth and promotes proliferation |
| 15 | NP_037481.1 | NENF | + | - | Increases cell proliferation and promotes hippocampal neurogenesis |

**Supplementary Table 2.** List of all the antibodies used in the study.

| Antibody | Company | Catalog No | Host | Application | Dilution |
|---|---|---|---|---|---|
| CHI3L1 | R&D Systems, Minneapolis, MN | AF2699 | Goat | IF | 1:200 |
| AQP1 | Santa Cruz Bioscience, Dallas, TX | sc-25287 | Rabbit | IF | 1:100 |
| Myocilin | Santa Cruz Bioscience Dallas, TX | | Rabbit | WB | 1:100 |
| Myocilin (55kDa) | Novus Biologicals, Littleton, CO | 297817 | Mouse | IF | 1:500 |
| ANGPTL7 | R&D Systems, Minneapolis, MN | MAB914 | Mouse | IF | 1:100 |
| COX2 (21kDa) | Santa Cruz Bioscience, Dallas, TX | sc-514489 | Mouse | IF/WB | 1:100/1:500 |
| GRP78 (78kDa) | Santa Cruz Bioscience Dallas, TX | sc-376768 | Mouse | WB | 1:500 |
| TMEM177(50kDa HC, 25kDa LC) | Invitrogen, Carlsbad, CA | A6-A11-9 | Mouse | IF/WB | 1:100/1:500 |
| β-Actin (50kDa) | Invitrogen, Carlsbad, CA | MA5-5739 | Mouse | WB | 1:5000 |
| CD90- BV510 | BD Bioscience, San Jose, CA | 563070 | Mouse | FC | 1:100 |
| CD73- PE/Cy7 | Biolegend, San Diego, CA | 344010 | Mouse | FC | 1:100 |
| CD105- AF647 | Biolegend, San Diego, CA | 323212 | Mouse | FC | 1:100 |
| CD166- FITC | MBL, Woburn, MA | K0044-4 | Mouse | FC | 1:100 |
| NOTCH1- PE | BD Pharmingen™, San Diego, CA | 563421 | Mouse | FC | 1:100 |
| OCT4-FITC | Santa Cruz Bioscience Dallas, TX | sc-5279 | Mouse | FC | 1:100 |
| SSEA4- AF488 | eBioscience Inc., Coraopolis, PA | 53-8843-42 | Mouse | FC | 1:100 |
| CD34- FITC | Millipore, Burlington, MA | CBL555F | Mouse | FC | 1:100 |
| CD45- PE | BD Bioscience, San Jose, CA | 553081 | Mouse | FC | 1:100 |
| ABCB5 | Abcam, Cambridge, MA | ab140667 | Mouse | IF | 1:100 |
| CD31 | BD Bioscience, San Jose, CA | 550274 | Rat | IF | 1:100 |
| Collagen IV | Sigma Aldrich, St. Louis, MO | SAB4500385 | Rabbit | IF | 1:100 |
| Fibronectin | Abcam, Cambridge, MA | Ab23750 | Rabbit | IF | 1:100 |
| OCT4 | Millipore, Burlington, MA | MAB4401 | Mouse | IF | 1:200 |
| Ki67 | Abcam, Cambridge, MA | ab15580 | Rabbit | IF | 1:50 |
| IgG1 K Iso FITC | Bioscience Inc. San Diego, CA | 11-4714-42 | Mouse | FC | 1:100 |
| IgG2a K Isotype PE | BD Pharmingen™, San Diego, CA | 555574 | Mouse | FC | 1:100 |
| IgG1 K Iso PE/CY7 | Biolegend, San Diego, CA | 400126 | Mouse | FC | 1:100 |
| IgG1 K Iso APC | Bioscience Inc. San Diego, CA | 17-4714-42 | Mouse | FC | 1:100 |
| Anti-goat AF-555 IgG | Life Technologies, Eugene, OR | A11056 | Donkey | IF | 1:1500 |
| Anti-rabbit AF-555 IgG | Life Technologies, Eugene, OR | A32794 | Donkey | IF | 1:1500 |
| Anti-mouse AF-647 IgG | Life Technologies, Eugene, OR | A31571 | Donkey | IF | 1:1500 |
| Anti-Rabbit AF-488 IgG | Invitrogen, Carlsbad, CA | A21206 | Donkey | IF | 1:1500 |
| Anti-Rat AF-488 IgG | Invitrogen, Carlsbad, CA | A21208 | Donkey | IF | 1:1500 |

**Abbreviations:** FITC-Fluorescein Isothiocyanate, PE-Phycoerythrin, APC- Allophycocyanin, HC-Heavy Chain, LC-Light Chain, IF-Immunofluorescence, WB-Western Blotting, FC-Flow Cytometry
